## Supplementary example on hypothetical data for "Considerations for using reproduction data in toxicokinetic-toxicodynamic modelling"

To illustrate the potential for bias in the parameter estimation, we will run through a few model fits with a hypothetical data set. The data set is constructed from real data from the control of a *Daphnia magna* reproduction study, for a single female. The aim of this exercise is to get an idea of the potential bias in a toxicity test with pulsed exposure. For the treatments, we use a two-day pulse with two arbitrary pulse heights (5 and 10 µg/L). The data for these treatments are copied from the control, but manipulated to correspond to a rapid and reversible effect on the costs for reproduction. The counts at $t$=15 are decreased by 50% relative to the control at 5 µg/L, and by 95% at 10 µg/L. We will use the complete data set, and a censored one in which the observations are removed where no reproduction was observed (see main text, Figure 2A-B). Only the initial zeros are left in, assuming a reproduction cycle of 3 days. Body length (mm) over time was based on the same *D. magna* test and taken the same in all treatments (since we assume an effect on reproduction costs). In the model fit, the initial body length was fixed to 0.924 mm.

*Table 1. Constructed data sets for cumulative reproduction over time (eggs or neonates).*

| | Original data set | | | | Censored data set | | |
|---|---|---|---|---|---|---|---|
| | 0 µg/L | 5 µg/L | 10 µg/L | | 0 µg/L | 5 µg/L | 10 µg/L |
| 0 | 0 | 0 | 0 | | 0 | 0 | 0 |
| 1 | 0 | 0 | 0 | | 0 | 0 | 0 |
| 2 | 0 | 0 | 0 | | 0 | 0 | 0 |
| 3 | 0 | 0 | 0 | | 0 | 0 | 0 |
| 4 | 0 | 0 | 0 | | 0 | 0 | 0 |
| 5 | 0 | 0 | 0 | | 0 | 0 | 0 |
| 6 | 0 | 0 | 0 | | 0 | 0 | 0 |
| 7 | 0 | 0 | 0 | | - | - | - |
| 8 | 0 | 0 | 0 | | - | - | - |
| 9 | 12 | 12 | 12 | | 12 | 12 | 12 |
| 10 | 12 | 12 | 12 | | - | - | - |
| 11 | 12 | 12 | 12 | | - | - | - |
| 12 | 41 | 41 | 41 | | 41 | 41 | 41 |
| 13 | 41 | 41 | 41 | | - | - | - |
| 14 | 41 | 41 | 41 | | - | - | - |
| 15 | 78 | 59 | 43 | | 78 | 59 | 43 |
| 16 | 78 | 59 | 43 | | - | - | - |
| 17 | 78 | 59 | 43 | | - | - | - |
| 18 | 114 | 95 | 79 | | 114 | 95 | 79 |
| 19 | 114 | 95 | 79 | | - | - | - |
| 20 | 114 | 95 | 79 | | - | - | - |
| 21 | 153 | 134 | 118 | | 153 | 134 | 118 |

*Table 2. Constructed data sets for body length (mm) over time.*

| | 0-10 μg/L |
|---|---|
| 0 | 0.924 |
| 7 | 2.908 |
| 14 | 3.577 |
| 21 | 3.874 |

We will use the same data set to show the impact of clutch-wise spawning and the impact of the brood-pouch delay. Rather than modifying the data set, we will shift the timing of the pulse exposure. For clutch-wise spawning, we shall assume that the data set is not for *D. magna*, but for a species where eggs are counted (so there is no delay caused by the incubation in the brood pouch). We place the pulse treatment at *t*=12-14, so the effect is observed at the following clutch of eggs at *t*=15. For the brood-pouch delay, we shall assume that the data set represents neonate counts, such as for *D. magna*. We then place the pulse treatment earlier, at *t*=9-11, so the effect is observed at the third brood at *t*=15. There can be no effect at the second brood (*t*=12) since the eggs for this brood were produced at *t*=9 and hence before the pulse exposure.

We use the simplified DEBkiss-based model (Jager 2020) and fit growth and reproduction simultaneously with the algorithm as described in Jager (2021). The likelihood function is based on the normal distribution, after square-root transformation. Confidence intervals are 95% likelihood-based intervals. The mode of action is set at costs for reproduction and no feedbacks for damage are included. In all cases, the basic DEB parameters are fitted to the control treatment first. Next, the basic parameters are fixed and only the toxicity-related parameters fitted to the complete data set. This is the strategy proposed by Jager (2020).

*Bias due to clutch-wise spawning*

Here, we assume that the counts for reproduction are eggs, so there is no brood pouch delay to worry about. The pulse is from *t*=12-14. The data are fitted to the complete data set (Figure 1A), and to the censored data set (Figure 1B). Fitting on the complete set shows that the control fit (Figure 1A-R0) represents an averaged-out version of the data set, as explained in the main text, since the model cannot reproduce the stepwise pattern in the data set. For the censored data set, the model curve is closer to all data points (Figure 1B-R0). Comparing the two control fits, there are small differences in all model parameters (Table 3), but the most striking difference is in the length at puberty: using the censored data set leads to a lower length at puberty, and thereby an earlier start of reproduction. The model fit on the original data set is biased towards a delayed reproduction because of the frequent 'zero-reproduction' observations in the data set.

The fits on the treatment data are also quite different. The censored data set (Figure 1B-R1/2) shows the rapid effect as intended during the construction of this data set (the only effect is a reduction of the eggs in a single clutch, produced at *t*=15). When fitting the original data set, however, the effect is smeared out over time (Figure 1A-R1/2): the dominant rate constant is somewhat lower, the threshold is lower, and the effect strength is much lower than for the censored data set (Table 3). This relates to the differences in effect patterns between the two fits.

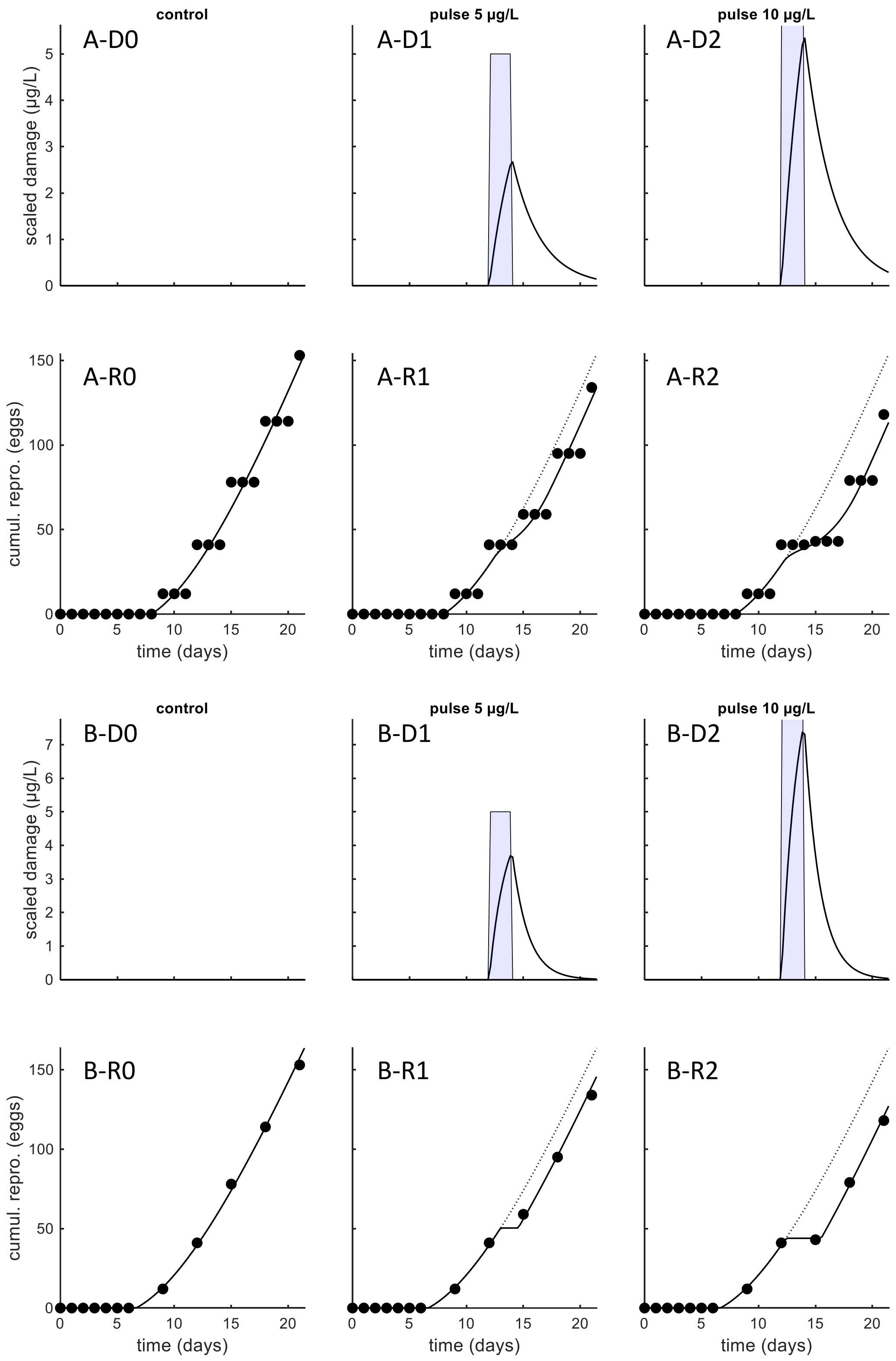


*Figure 1. Fits of the DEB-TKTD model on the artificial data for egg-clutch production. The top panels (A-D and A-R) show the fits on the original data set, and the lower panels (B-D and B-R) the fit on the censored data set (observations with zero-reproduction are removed).*

*Table 3. Results of the parameter estimation for the various cases. Asterisk indicates that the estimate or its confidence interval have reached the maximum or minimum value allowed in the optimisation.*

| | **Counts are eggs pulse treatment at *t*=12-14** | | **Counts are neonates pulse treatment at *t*=9-11** | | |
|---|---|---|---|---|---|
| Analysis approach | Original data No BP shift | Censored data No BP shift | Original data No BP shift | Censored data No BP shift | Censored data 3-day BP shift in model |
| Fits shown in … | Figure 1A | Figure 1B | Figure 2 | Figure 3A | Figure 3B |
| | **Fit basic parameters on control data** | | | | |
| Length at puberty ($L_p$, mm) | 3.05 (2.98-3.10) | 2.84 (2.78-2.91) | 3.05 (2.98-3.10) | 2.84 (2.78-2.91) | 2.06 (1.99-2.18) |
| Maximum length ($L_m$, mm) | 4.00 (3.94-4.06) | 3.99 (3.91-4.05) | 4.00 (3.94-4.06) | 3.99 (3.91-4.05) | 3.99 (3.91-4.05) |
| Von B. growth rate constant ($r_B$, 1/d) | 0.147 (0.139-0.156) | 0.148 (0.140-0.161) | 0.147 (0.139-0.156) | 0.148 (0.140-0.161) | 0.148 (0.140-0.160) |
| Max. reproduction rate ($R_m$, eggs/d) | 17.6 (16.0-19.0) | 16.7 (16.0-17.5) | 17.6 (16.0-19.0) | 16.7 (16.0-17.5) | 17.6 (16.6-18.6) |
| | **Fit tox parameters on all data (basic parameters fixed)** | | | | |
| Dominant rate constant ($k_d$, 1/d) | 0.397 (0.177-0.898) | 0.724 (0.608-0.885) | 0.01* (0.01*-0.135) | 0.0857 (0.01*-0.188) | 0.694 (0.555-0.892) |
| Threshold for effects ($z_b$, µg/L) | 0.736 (0.000757*-2.11) | 2.57 (2.03-2.76) | 0.0267 (0.000757*-0.340) | 0.188 (0.000757*-0.421) | 2.56 (1.59-2.79) |
| Effect strength ($b_b$, L/µg/d) | 0.693 (0.281-7.44) | 1040* (4.90-1040*) | 3.36 (0.308-5.84) | 0.579 (0.292-4.22) | 1040* (2.14-1040*) |

### *Bias due to brood-pouch delay*

Here, we assume that the counts for reproduction are released neonates, so this is representative for *D. magna* with brood pouch. The pulse is now from *t*=9-11; the first egg clutch that is produced after the pulse is at *t*=12, and is released at *t*=15.

We start with the fit for the original data set when we ignore both clutch-wise spawning (no removal of data points) and brood-pouch delay (no shift in the model). Note that the control fit without shift (Figure 2-R0) is the same as for the egg-count fit on the original data set of the previous section (Figure 1A-R0). Small differences would be possible since the optimisation algorithm contains a random element.

The parameter estimates are included in Table 3 above. Using the original data set without shift provides a huge distortion of the effect pattern (Figure 2-R1/2). The short-lasting and rapid effect on reproductive investment is turned into a very gradual and prolonged change in reproduction rate in the model fit. This is also reflected in the parameter estimates: the dominant rate constant goes to its lowest allowed value in the optimisation (i.e., ‘slow kinetics’).

In Figure 3, the fits to the censored data set are shown. The first fit (Figure 3A) ignores the delay caused by the brood pouch, while the second fit (Figure 3B) shifts the model prediction for reproduction by 3 days. Note that the control fit without shift (Figure 3A-R0) is the same as for the egg-count fit on the censored data set of the previous section (Figure 1B-R0). Including the shift leads to a substantial reduction in the length at puberty (Table 3). Puberty is now a better estimate of the point where investment in reproduction starts, which is before the production of the first clutch of eggs, which in turn is well before the release of the first batch of neonates.

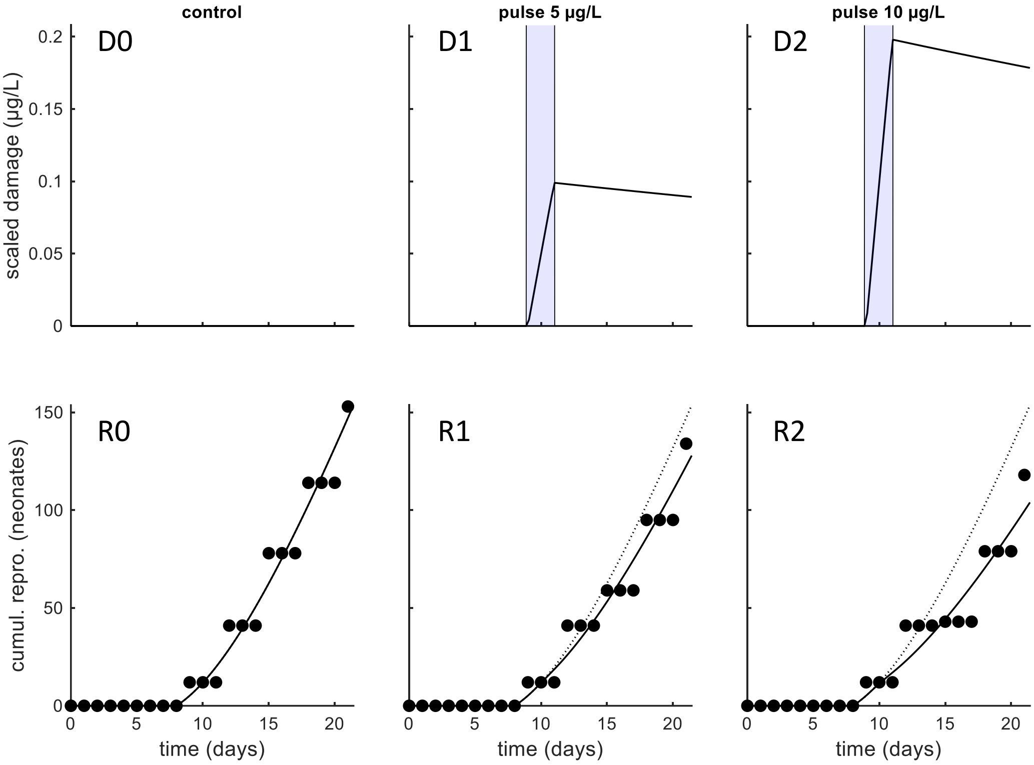


*Figure 2. Fits of the DEB-TKTD model on the artificial data for neonate counts. The panels show the fits without a brood-pouch delay on the original data set (no censoring) and without a brood-pouch delay.*

The fit when ignoring the brood-pouch delay (Figure 3A-R1/2) shows very slow damage dynamics, and a very gradual effect over time. The model is not able to capture the stop in reproduction in the high treatment at all (Figure 3A-R2). The reason is that the effect occurs *after* the pulse has already ended. Since the effect is on the reproductive investment, and the brood-pouch delay is ignored, the model is unable to properly capture the pattern in the data set. With the brood-pouch delay accounted for in the model, the fit is as expected (Figure 3B-R1/2), and the parameter estimates are actually almost identical to the ones for the clutch-wise spawning on the censored data set (Table 3).

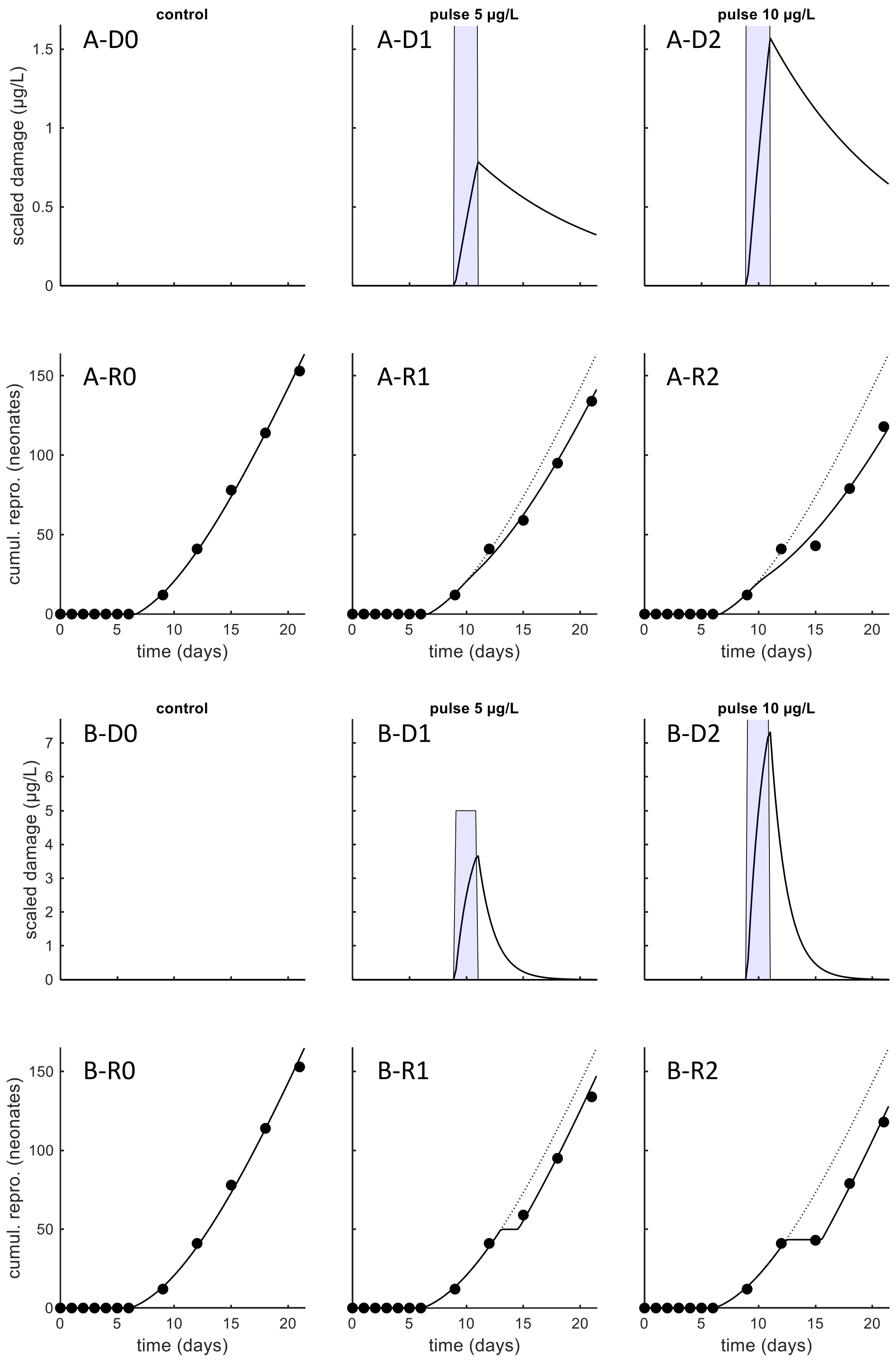


*Figure 3. Fits of the DEB-TKTD model on the artificial data for neonate counts. Only the censored data set is used. The top panels (A-D and A-R) show the fits without a brood-pouch delay, and the lower panels (B-D and B-R) the fit of the model with a 3-day brood pouch delay included.*

*Concluding on this example*

This is just a single, perhaps rather arbitrary, example, for a particular life history, particular exposure pattern, and for a chemical with specific properties. Therefore, no general conclusions can be drawn, other than that ignoring the issues of clutch-wise spawning and brood-pouch delay can indeed lead to substantial bias in the model fit and the model parameters. This example also shows that censoring the data set, and shifting the model prediction, is a simple and effective way to obtain more realistic model fits from the data set.

Bias in the model parameters will lead to bias in model predictions. The extent of this prediction bias will depend on the chemical, the species, the test conditions, but will also strongly depend on the exposure scenario to which we are extrapolating. This example shows that for a chemical with rapid damage dynamics, we should expect that ignoring clutch-wise spawning and brood-pouch delays leads to lower values for the dominant rate constant $k_d$, thus giving the appearance of slower damage dynamics. The most extreme situation is shown in Figure 2-D1/2, where the damage level hardly decreases after the pulse (compare this to the damage patterns in Figure 3B-D1/2). As a result, ignoring these issues will overestimate carry-over toxicity (Ashauer et al. 2010), where subsequent pulses will lead to an increase of the toxic effect. Furthermore, toxicity will be overestimated for prolonged or constant exposure, as damage will build up over longer exposure. However, for chemicals with slow damage dynamics, a different bias might result.
